# Expanding the SiMPl plasmid toolbox for use with spectinomycin/streptomycin

**DOI:** 10.1101/2021.02.04.429595

**Authors:** Navaneethan Palanisamy, Jara Ballestin Ballestin, Barbara Di Ventura

## Abstract

We recently developed the SiMPl plasmid toolbox, which is constituted by pairs of plasmids, generically indicated as pSiMPl^x^_N and pSiMPl^x^_C, which can be stably maintained in *Escherichia coli* with a single antibiotic x. The method exploits the split intein gp41-1 to reconstitute the enzyme conferring resistance towards the antibiotic x, whereby each enzyme fragment is expressed from one of the plasmids in the pair. pSiMPl plasmids are currently available for use with ampicillin, kanamycin, chloramphenicol, hygromycin and puromycin. Here we introduce another pair for use with spectinomycin/streptomycin broadening the application spectrum of the SiMPl toolbox. To find functional splice sites in aminoglycoside adenylyltransferase we apply a streamlined strategy looking exclusively at the flexibility of native cysteine and serine residues, which we first validated splitting the enzymes conferring resistance towards ampicillin, kanamycin, chloramphenicol and hygromycin. This strategy could be used in the future to split other enzymes conferring resistance towards antibiotics.

---

We have recently devised a method based on <u>s</u>plit <u>i</u>ntein-<u>m</u>ediated enzyme reconstitution, that allows selecting bacterial and mammalian cells containing two <u>p</u>lasmids with a single antibiotic, which we named SiMPl^*1*^. Inteins are proteins that excise themselves out of host proteins in an autocatalytic reaction that results with a peptide bond joining the polypeptides originally flanking the intein^*2, 3*^. Split inteins are constituted by two separate fragments that need to associate to reconstitute a functional intein. Therefore, during the process of splicing, split inteins make fusions between two previously independent proteins or peptides. The SiMPl method consists in splitting the enzyme conferring resistance towards the antibiotic of choice – here indicated as x – in an N- and a C-terminal fragment, each *per se* dysfunctional. The coding sequence of the N-terminal fragment of the enzyme is fused to the coding sequence of the N-terminal fragment of the gp41-1 split intein^*4*^ and is cloned in one plasmid in place of the full-length resistance gene, giving rise to the pSiMPl^x^_N plasmid. The coding sequence of the C-terminal fragment of the enzyme is likewise fused to the coding sequence of the C-terminal fragment of gp41-1 and is cloned in another plasmid, giving rise to the pSiMPl^x^_C plasmid. The pSiMPl^x^_N and pSiMPl^x^_C plasmid pair for use in bacteria is characterized by compatible origins of replication and can be co-transformed into competent bacterial cells. Those cells that pick both plasmids up following the transformation procedure survive in the presence of the antibiotic x because the full-length enzyme that confirms resistance towards x gets reconstituted inside them. Thus, these cells are able to form colonies on the selection plate. SiMPl is, as the same says, extremely simple in concept and usage. The challenge consisted in identifying functional splice sites in the enzymes conferring resistance towards the antibiotics. To this end, we applied a rational strategy consisting in looking at (i) the flexibility of the amino acids ––using the coarse-grained CABS-flex protein structure flexibility simulation tool^*5*^ that calculates the fluctuations of the Cα atoms (root mean square fluctuation or RMSF) of the protein backbone structural ensemble––, (ii) their conservation, (iii) their surface accessibility and (iv) their location relative to the active site^*1*^. We considered all these features because, to ensure high splicing efficiency, we decided to insert in the enzymes, beyond the catalytic serine, five additional amino acids, called local exteins (SGYSS), which are residues found in the natural context of gp41-1^*4*^. However, when looking for additional splice sites in puromycin acetyltransferase (PAT) that would give rise to differently-sized enzyme fragments, we realized that identifying highly flexible native serines or cysteines also led to functional splice sites^*1*^. The local exteins appeared not to be necessary, and gp41-1 was able to splice itself out and reconnect the flanking enzyme fragments using either serine or cysteine as catalytic residue to a degree compatible with the growth of the double transformants in the presence of puromycin. Scarless reconstitution has the advantage that the activity of the reconstituted enzyme is most likely the same as that of the full-length, native enzyme. Here we wished to assess whether this streamlined strategy looking exclusively at the flexibility of native serines and cysteines could be generally applied to enzymes conferring resistance towards antibiotics. We first validate the method identifying many new functional splice sites for the enzymes conferring resistance towards ampicillin, kanamycin, chloramphenicol and hygromycin. Then, using this strategy, we find functional splice sites for aminoglycoside adenylyltransferase and establish a novel SiMPl plasmid pair (pSiMPl^s^) for use with spectinomycin/streptomycin.

## RESULTS AND DISCUSSION

To validate whether a streamlined strategy, whereby only the flexibility of native serines and cysteines is considered, would lead to the prediction of functional splice sites on enzymes conferring resistance towards antibiotics, we first considered the same enzymes we previously split: aminoglycoside 3′-phosphotransferase, the enzyme conferring resistance towards kanamycin (for simplicity, in the following referred to as APT); chloramphenicol acetyltransferase, the enzyme conferring resistance towards chloramphenicol (in the following referred to as CAT); TEM-1 beta-lactamase, the enzyme conferring resistance towards ampicillin (in the following referred to as TEM-1 β-L); and hygromycin B phosphotransferase, the enzyme conferring resistance towards hygromycin (in the following referred to as HPT). For all these enzymes, the crystal structure is available. Using CABS-flex^*5*^, we calculated the flexibility of Cα of each amino acid (given as RMSF) and took the serines and cysteines for which the RMSF was highest (**Figure 1a,c,e,g)**. We predicted six, five, four and seven splice sites for APT, CAT, TEM-1 β-L and HPT, respectively (**Figure 1a,c,e,g)**. We cloned the corresponding N- and C-terminal fragments of each enzyme in the previously constructed pSiMPl_N/C plasmid pairs, respectively, and then transformed them either individually or together in *E. coli* TOP10 cells. pSiMPl_N plasmids carry the *egfp* gene cloned in the multiple cloning site, while pSiMPl_C ones carry the *mruby3* gene. For most predicted splice sites, we obtained colonies only when co-transforming both pSiMPl plasmids (**Figure 1b,d,f,h and Fig. S1)**. One of the six predicted splice sites for APT (Q35:S36) and one of the five predicted splice sites for CAT (I13:S14) resulted in the C-terminal fragment of the enzymes having some level of activity (**Figure 1b** and **1d**). Similarly, for TEM-1 β-L, splice sites R220:S221 and G263:S264 resulted in N-terminal fragments having some activity (**Figure 1f**). When using information solely on structure flexibility, it is indeed impossible to computationally predict whether the fragments will have activity on their own or not. However, it is easy to experimentally identify such cases and exclude them afterwards. Recently, Ho and colleagues proposed an empirical method to identify functional splice sites on proteins of interest called intein-assisted bisection mapping (IBM)^*6*^. With this method, the authors predicted two hotspots for splice sites in TEM-1 β-L; splice site G263:S264 predicted by our computational method lies in the second hotspot identified by IBM^*6*^. However, our experimental results showed that the N-terminal fragment of the enzyme split at this position has some activity, thus we would not consider this a functional site. Likewise, Jillette and colleagues applied a purely experimental pipeline based on trial-and-error and successfully identified eight splice sites for HPT^*7*^ (tested in mammalian cells), among which three found by our rational approach as well. This indicates that several strategies exist to find functional splice sites in enzymes conferring resistance towards antibiotics. We note that being ours a purely computational method, it is less cumbersome than any other purely experimental one, as it restricts the numbers of samples to test beforehand.

**Figure 1.**
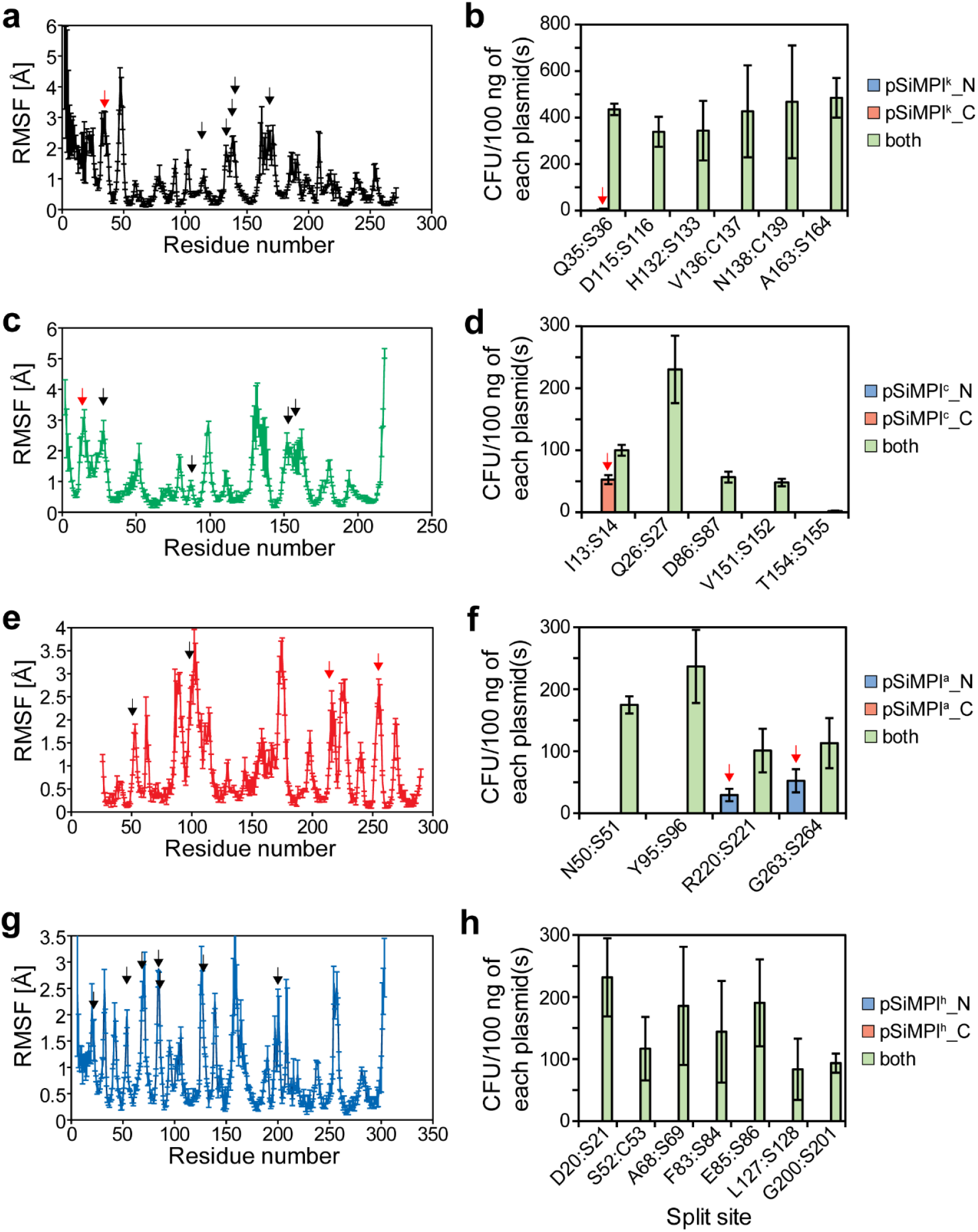
Identification of splice sites in enzymes conferring resistance towards antibiotics using a streamlined computational strategy. (**a,c,e,g**) Root mean square fluctuation (RMSF) of Cα atoms in aminoglycoside 3′-phosphotransferase (PDB ID: 4ej7; **a**), chloramphenicol acetyltransferase (PDB ID: 1q23; **c**), TEM-1 β-lactamase (PDB ID: 1zg4; **e**) and hygromycin B phosphotransferase (PDB ID: 3w0s; **g**) obtained by submitting the crystal structures to the CABS-flex 2.0 web-server*5*. Values represent mean ± S.E.M. of three independent simulations with different seed values. Black arrow, functional splice site. Red arrow, splice site corresponding to one of the two enzyme fragments having activity. (**b,d,f,h**) Bar graph showing the transformation efficiency of the indicated SiMPl plasmids in chemically competent *E. coli* TOP10 cells grown on kanamycin (**b**), chloramphenicol (**d**), ampicillin (**f**) and hygromycin (**h**) selective plates after Gibson Assembly®. Values represent mean ± S.E.M. of three independent experiments. Red arrow, splice site corresponding to one of the two enzyme fragments having activity.

All splice sites found to be functional were surface exposed (**Figures S2** and **S3**), suggesting that high flexibility largely reflects accessibility as well. Agarose gel electrophoretic analysis of the plasmid DNA extracted from a randomly-picked colony indicated the presence of both pSiMPl plasmids for all antibiotics tested (**Figure S4**). Moreover, polymerase chain reaction (PCR) confirmed the presence of the genes of interest (*egfp* and *mruby3*) (**Figures S5 and S6**), and the absence of an intact resistance gene (**Figure S7**). We also verified by Western blot that protein splicing occurred in all cases (**Figures S8-S11**).

Next, we characterized the different splice sites in terms of bacterial growth and transformation efficiency. *E. coli* TOP10 cells were transformed with all pSiMPl plasmid pairs corresponding to the different splice sites of the various enzymes and cultures were grown in 96-well microtiter plates in nutrient broth with the appropriate antibiotics (**Figure S12 a-d**). All splice sites supported bacterial growth in liquid culture to a similar extent, with few exceptions showing a longer lag phase (splice site T154:S155 for CAT, and splice sites F83:S84 and L127:S128 for HPT). This was also reflected in the generation time, which was similar for most constructs (**Figure S12e**). Since TEM-1 β-L also confers resistance towards carbenicillin (a variant of ampicillin; **Figure S13a,b**), we tested the two newly identified splice sites as well as the one previously identified (S104:P105^*1*^) with this antibiotic and found that they all supported bacterial growth to the same extent as in the presence of ampicillin (**Figure S13d-f**). As positive control we took the full-length enzyme encoded on pTrc99a (**Figure S13c**). The generation time for bacteria expressing the newly identified split enzyme variants was indistinguishable for media containing ampicillin and carbenicillin (**Figure S13g**). The transformation efficiency for the pSiMPl pairs for usage with kanamycin, chloramphenicol and ampicillin was equivalent to that of conventional, full-length enzyme-bearing plasmids only at high total plasmid concentration (100 ng; **Figure S14**). At lower total plasmid concentrations (1 and 0.1 ng), on the other hand, the transformation efficiencies significantly decreased and varied between the different constructs (**Figure S14**). It is to be noted, though, that we are comparing transformation of the pSiMPl pairs to transformation of single plasmids.

We previously observed the spontaneous appearance of a mutation within reconstituted HPT split at E255:L256, which was likely necessary for the function of the enzyme^*1*^. Interestingly, also for some of the new split versions of HPT, a mutation within the N-terminal fragment was acquired (**Figure S15a**). In some cases, instead of a mutation within the coding sequence, an insertion within the promoter arose (**Figure S15b**). We speculate that these mutations have the effect of increasing the enzyme activity and expression level, respectively, thus bacteria carrying them would have an advantage over the other ones. We indirectly tested this hypothesis by comparing the growth curves and generation times of bacteria carrying either wild-type HPT, HPT^T9R^ or the construct with the mutated promoter. The advantage of the mutations was particularly evident through the generation time, which was shorter for both mutations than for the wild type (**Figure S15c,d**).

Finally, we checked whether the streamlined strategy could be applied with another split intein, namely *Npu* DnaE, which has cysteine as catalytic residue^*8*^. We took the splice sites we already identified when using gp41-1 in correspondence to native cysteines–namely V136:C137 and N138:C139 in APT and S52:C53 in HPT –and tested if they supported bacterial growth when *Npu* DnaE was used instead of gp41-1. We obtained colonies only for splice site V136:C137 in APT. The splice site in HPT gave also rise to colonies. While for the APT construct based on the splice site V136:C137 there was no difference between *Npu* DnaE- or gp41-1-mediated enzyme reconstitution in terms of growth in liquid medium (**Figure S16a,b**), for the HPT construct based on splice site S52:C53, bacterial growth was better with gp41-1 than *Npu* DnaE (**Figure S16c,d**). These data suggest that the efficiency of splicing and, consequently, the amount of reconstituted HPT be higher when using gp41-1. This is interesting and at first counterintuitive, considering that cysteine is the natural catalytic residue for *Npu* DnaE and not gp41-1. Nonetheless, gp41-1 is the fastest split intein known to date, thus we speculate that the slightly suboptimal splicing due to the presence of the cysteine instead of the serine at the catalytic site results nonetheless in more efficient splicing than the one achieved by *Npu* DnaE.

After validating the streamlined strategy to locate splice sites using APT, CAT, TEM-1 β-L and HPT, we proceeded with its application to aminoglycoside adenylyltransferase, the enzyme that confers resistance towards spectinomycin/streptomycin. First, we needed to model the 3D structure of the enzyme. We used the fully automated protein structure homology-modelling server SWISS-MODEL^*9*^ with the crystal structure of *Salmonella enterica* AadA as template (PDB ID: 5g4a). Using again CABS-flex^*5*^ on the model structure, we calculated the flexibility of Cα of each amino acid (given as RMSF) and identified five serines for which the RMSF was highest (**Figure 2a)**. We then constructed pSiMPl^s^_N and pSiMPl^s^_C for all predicted splice sites, and found them to be functional (**Figure 2b**). Splice site T73:S74 led to fewer colony forming units than the other sites. Interestingly, we noticed that splitting chloramphenicol acetyltransferase at T154:S155 also led to fewer colonies compared to the other split versions. Whether splitting between a threonine and a serine is always problematic is worth future exploration. As before, we confirmed that the splice sites were surface exposed (**Figure S17**) and that both pSiMPl plasmids were present in the bacteria (**Figure 2c**). PCR confirmed the presence of the *egfp* and *mruby3* genes, and the absence of the full-length resistance gene (**Figure 2d**). Finally, we measured the growth curves of *E. coli* TOP10 cells transformed with all the pSiMPl^s^ plasmid pairs on spectinomycin (**Figure 2e**) and streptomycin (**Figure 2g**). Despite not being all identical, there weren’t major differences in the growth curves for the different splice sites. The generation times were also minimally lower for four out of the five constructs compared to the full-length enzyme for both antibiotics (**Figure 2f,h**).

**Figure 2.**
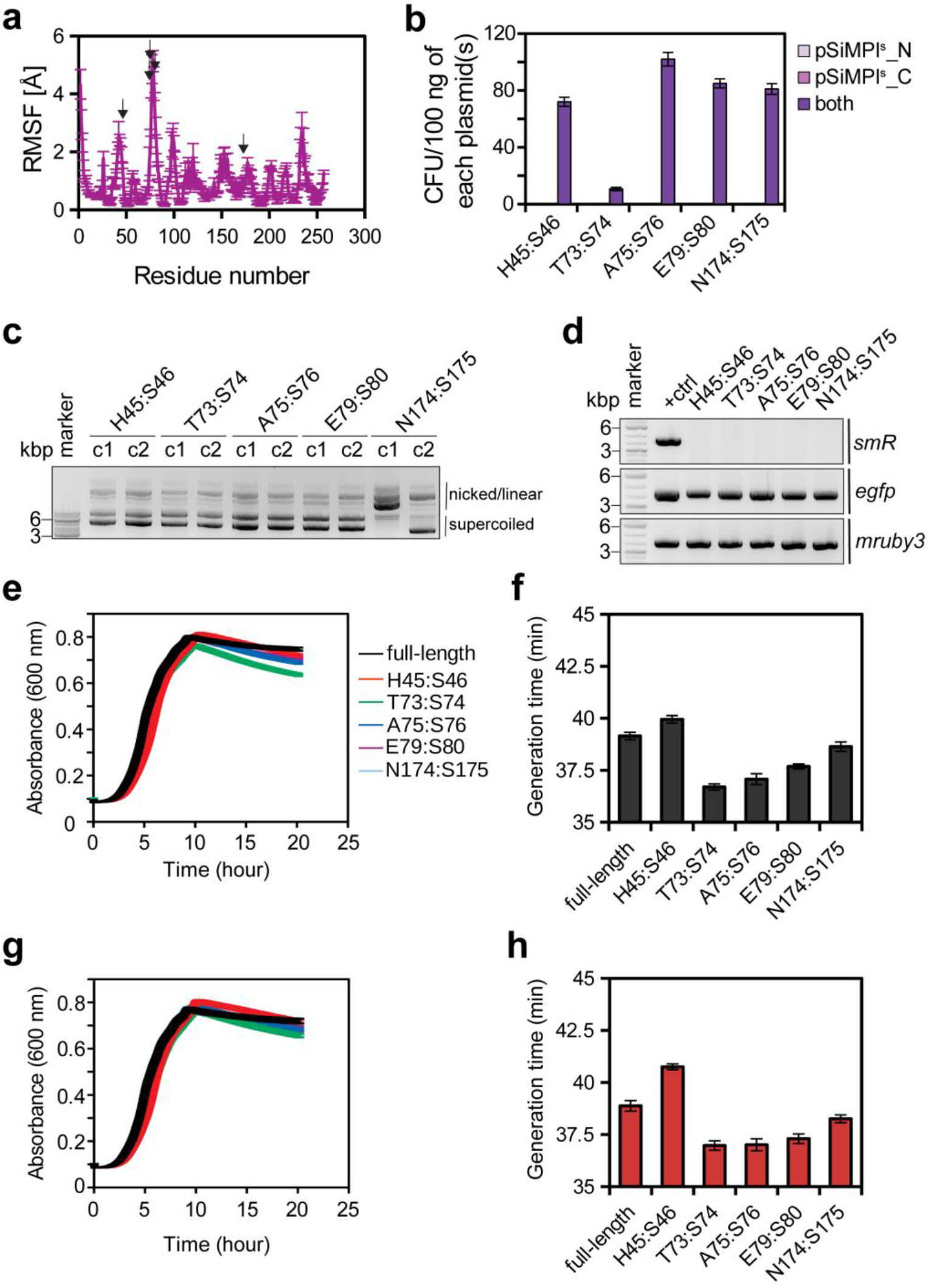
Identification of splice sites in aminoglycoside adenylyltransferase. (**a**) Root mean square fluctuation (RMSF) of Cα atoms in aminoglycoside adenylyltransferase obtained by submitting to the CABS-flex 2.0 web-server^*5*^ the 3D model structure obtained by SWISS-MODEL^*9*^ using the crystal structure of *Salmonella enterica* AadA as template (PDB ID: 5g4a). Values represent mean ± S.E.M. of three independent simulations with different seed values. Black arrow, functional splice site. (**b**) Bar graph showing the transformation efficiency of the indicated plasmids in chemically competent *E. coli* TOP10 cells grown on spectinomycin after Gibson Assembly®. Values represent mean ± S.E.M. of three independent experiments. (**c**) Ethidium bromide-stained agarose gel showing plasmid DNA isolated from a randomly-picked clone obtained after transformation with the SiMPls plasmids for selection on spectinomycin/streptomycin. (**d**) PCR analysis of the SiMPls plasmids isolated from bacteria. pBAD33 into which the full-length *smR* resistance gene (derived from pCDF-1b) was cloned in place of the typical *camR* was used as control to show the product obtained after amplification of the full-length resistance gene. (**e,g**) Growth curve of *E. coli* TOP10 cells transformed with the pSiMPls plasmids and grown in a 96-well plate in a medium containing spectinomycin (**e**) or streptomycin (**g**) measured every 2.4 min for 20 hours in a plate reader. Experiments were performed twice (biological replicates) with technical triplicates each time. Values represent mean ± S.E.M. (**f**,**h**) Bar graph showing the generation time for the growth curves shown in (**e**) and (**g**), respectively, calculated using the R package Growthcurver^*12*^.

Here we presented a very simple strategy to identify splice sites in enzymes conferring resistance towards antibiotics, which led to a success rate of 100%. However, we believe that, despite being an important feature, structure flexibility alone would not suffice to find, within a generic protein of interest, splice sites corresponding to a sufficient activity with high success rate. Indeed, previously, structure flexibility was found not to be a good feature to predict functional splice sites in proteins^*10*^. A currently available computational method to predict split sites on any protein of interest, indeed incorporates several other features beyond structure flexibility^*11*^. Nonetheless, this easy strategy seems to be well-suited for the specific case in which the screening assay is connected to survival.

## Author Contributions

NP conceived the study, performed computational analyses, and all cloning and bacterial experiments. JBB performed Western blotting. BDV supervised the work, secured funding and wrote the manuscript with input from NP and JBB.

## Notes

The authors declare no competing financial interest. The pSiMPl^s^ plasmids will be deposited on Addgene.

## ACKNOWLEDGMENTS

We thank Mehmet Ali Öztürk for discussions and critical reading of the manuscript. This work was funded by the Deutsche Forschungsgemeinschaft (DFG) under Germany’s Excellence Strategy through EXC294 (BIOSS—Center for Biological Signalling Studies) and EXC2189 (CIBSS—Centre for Integrative Biological Signalling Studies, Project ID 390939984).

## Notes

### Competing Interest Statement

The authors have declared no competing interest.

