## Supplementary Materials for "Expanding the SiMPl plasmid toolbox for use with spectinomycin/streptomycin"

### Supporting Information

Content

**Supplementary Figures S1-S17**
**Materials and methods**
**References**

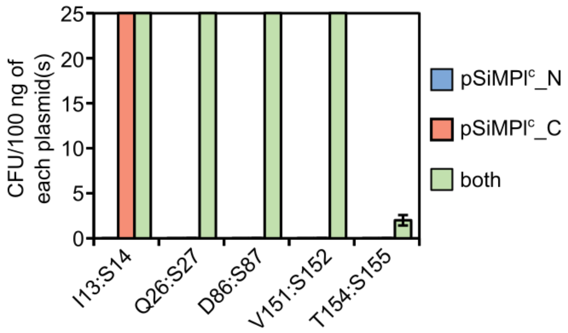


**Figure S1. Chloramphenicol acetyltransferase split at T154:S155 is functional.** Zoom in of the bar graph shown in **Figure 1d**. The transformation efficiency of the pSiMPl$^{c}$ plasmids in chemically competent *E. coli* TOP10 cells was assessed on chloramphenicol selective plates after Gibson Assembly®. Values represent mean ± S.E.M. of three independent experiments.

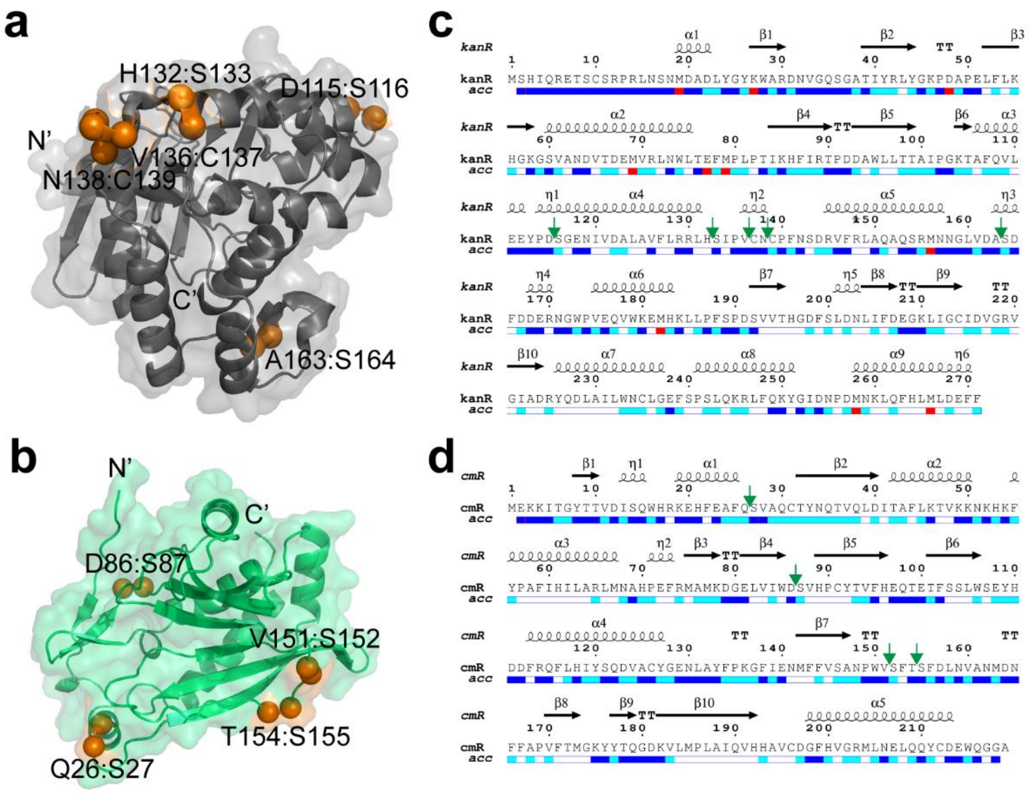


**Figure S2. The identified splice sites in aminoglycoside 3′-phosphotransferase and chloramphenicol acetyltransferase are surface accessible.** (**a**-**b**) Cartoon-surface representation of the crystal structure of aminoglycoside 3′-phosphotransferase (PDB ID: 4ej7; **a**) and chloramphenicol acetyltransferase (PDB ID: 1q23; **b**). The residues at the splice site are indicated by orange spheres. Structures were depicted in PyMOL (PyMOL Molecular Graphics System, v. 1.8.x, Schrödinger, LLC). (**c**-**d**) Relative accessibility (acc) of each residue in aminoglycoside 3′-phosphotransferase (**c**) and chloramphenicol acetyltransferase (**d**). Secondary structural elements are represented by black arrows and coils. The image was created using the ESPript 3.0 server[1]. The color code is as follow: blue = accessible, cyan = partially accessible, white = buried, red = not predicted. Splice site are indicated by green arrows.

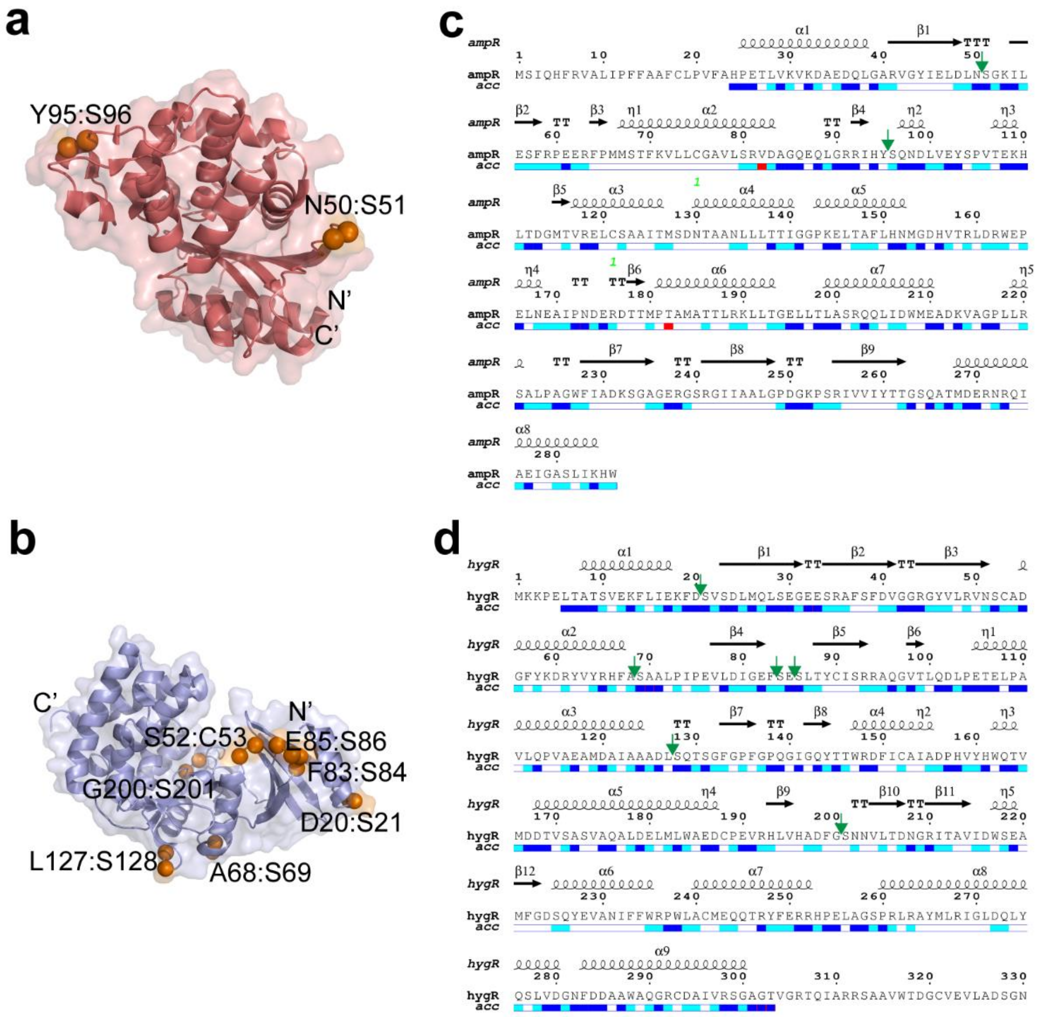


**Figure S3. The identified splice sites in TEM-1 β-lactamase and hygromycin B phosphotransferase are surface accessible.** (**a**-**b**) Cartoon-surface representation of the crystal structure of TEM-1 β-lactamase (PDB ID: 1zg4; **a**) and hygromycin B phosphotransferase (PDB ID: 3w0s; **b**). The residues at the splice site are indicated by orange spheres. Structures were depicted in PyMOL (PyMOL Molecular Graphics System, v. 1.8.x, Schrödinger, LLC). (**c**-**d**) Relative accessibility (acc) of each residue in TEM-1 β-lactamase (**c**) and hygromycin B phosphotransferase (**d**). Secondary structural elements are represented by black arrows and coils. The image was created using the ESPript 3.0 server[1]. The color code is as follow: blue = accessible, cyan = partially accessible, white = buried, red = not predicted. Splice site are indicated by green arrows.

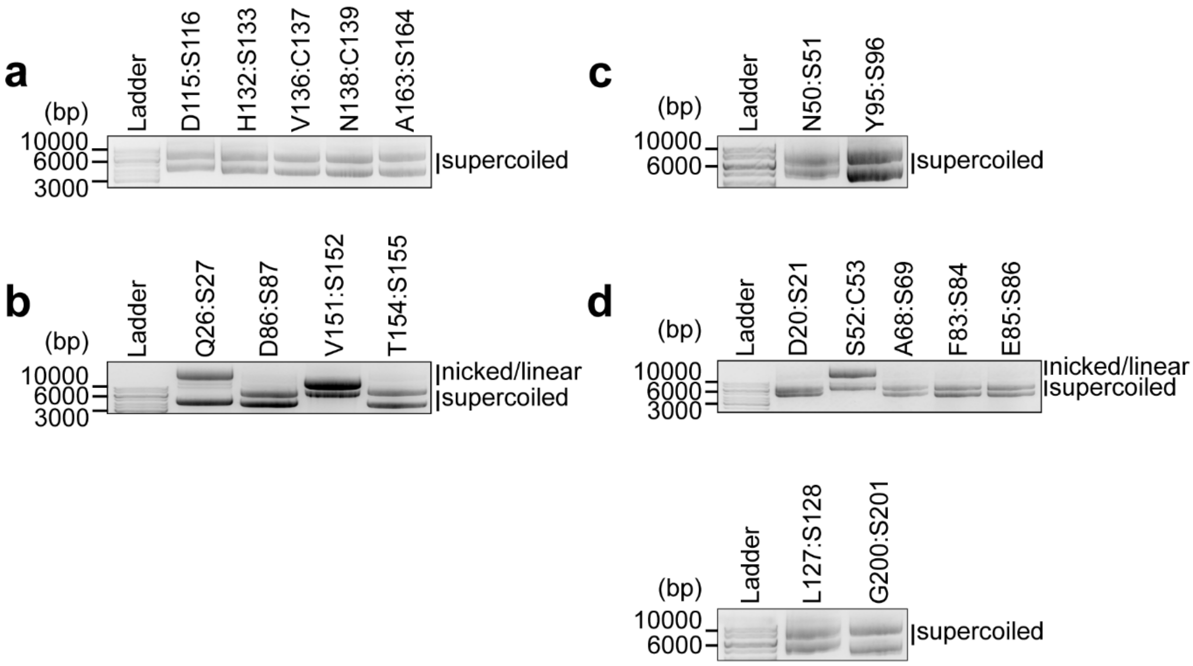


**Figure S4. Both pSiMPl plasmids are present in the cells.** (**a**-**d**) Representative ethidium bromide-stained 1% agarose gel showing that both plasmids are recovered after plasmid DNA extraction from a random bacterial clone. Cells were transformed with the pSiMPl plasmid pair for use with kanamycin (**a**), chloramphenicol (**b**), ampicillin (**c**) and hygromycin (**d**).

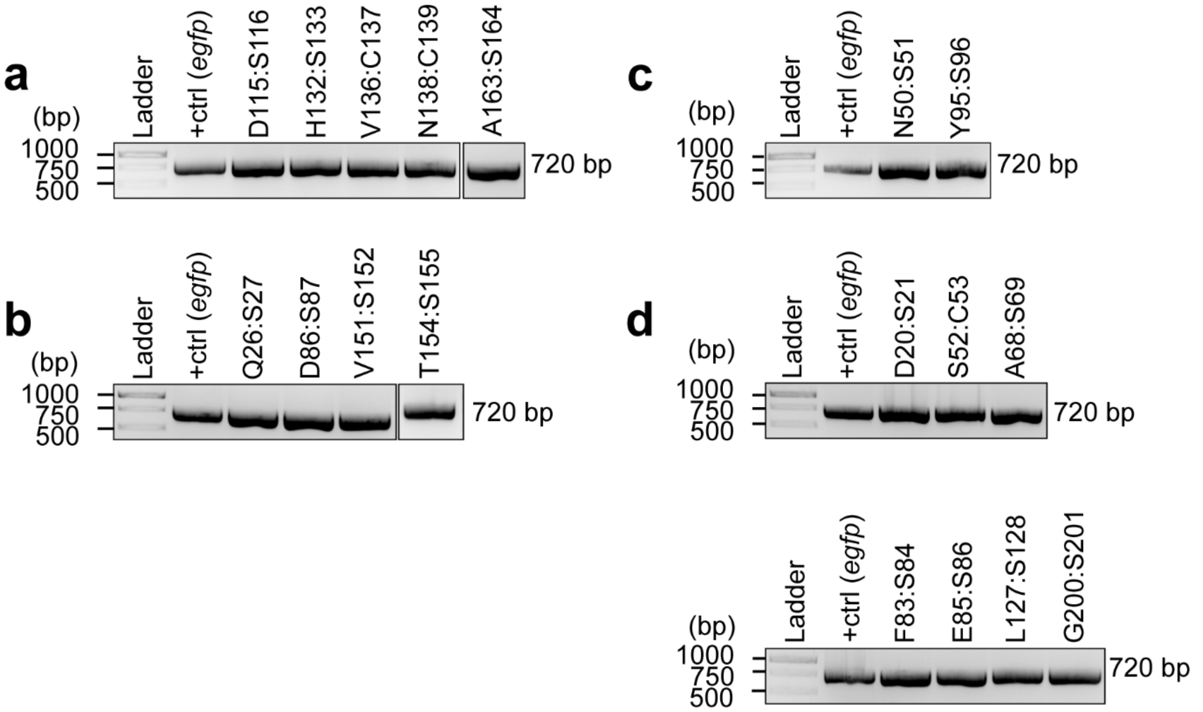


**Figure S5. The *egfp* gene encoded on pSiMPl_N is present in the cells transformed with pSiMPl plasmid pairs.** (**a**-**d**) Representative ethidium bromide-stained 1% agarose gel showing the PCR product obtained with *egfp*-specific primers. The PCR was performed on the plasmid DNA extracted from a random clone of cells transformed with the pSiMPl pair for use with kanamycin (**a**), chloramphenicol (**b**), ampicillin (**c**) and hygromycin (**d**). *egfp* cloned into pBAD33 was used as positive control (+ crtl).

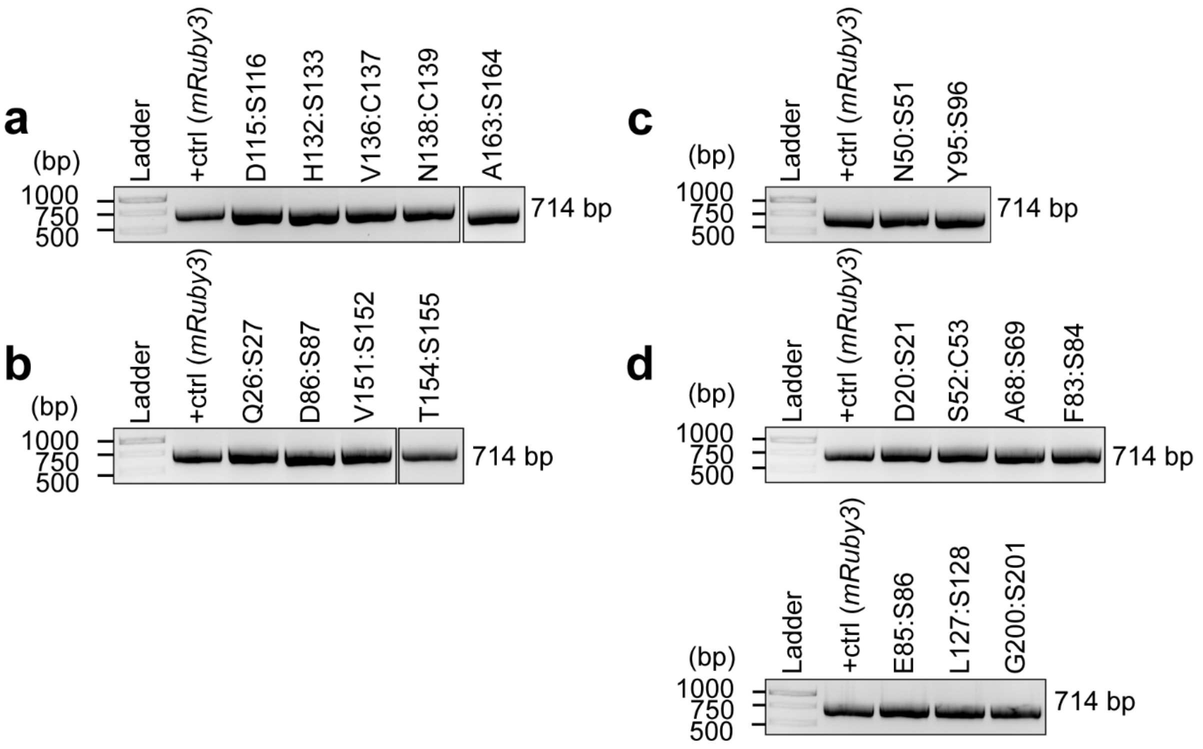


**Figure S6. The *mruby3* gene encoded on pSiMPl_C is present in the cells transformed with pSiMPl plasmid pairs.** (**a**-**d**) Representative ethidium bromide-stained 1% agarose gel showing the PCR product obtained with *mruby3*-specific primers. The PCR was performed on the plasmid DNA extracted from a random clone of cells transformed with the pSiMPl pair for use with kanamycin (**a**), chloramphenicol (**b**), ampicillin (**c**) and hygromycin (**d**). *mruby3* cloned into pTrc99a was used as positive control (+ crtl).

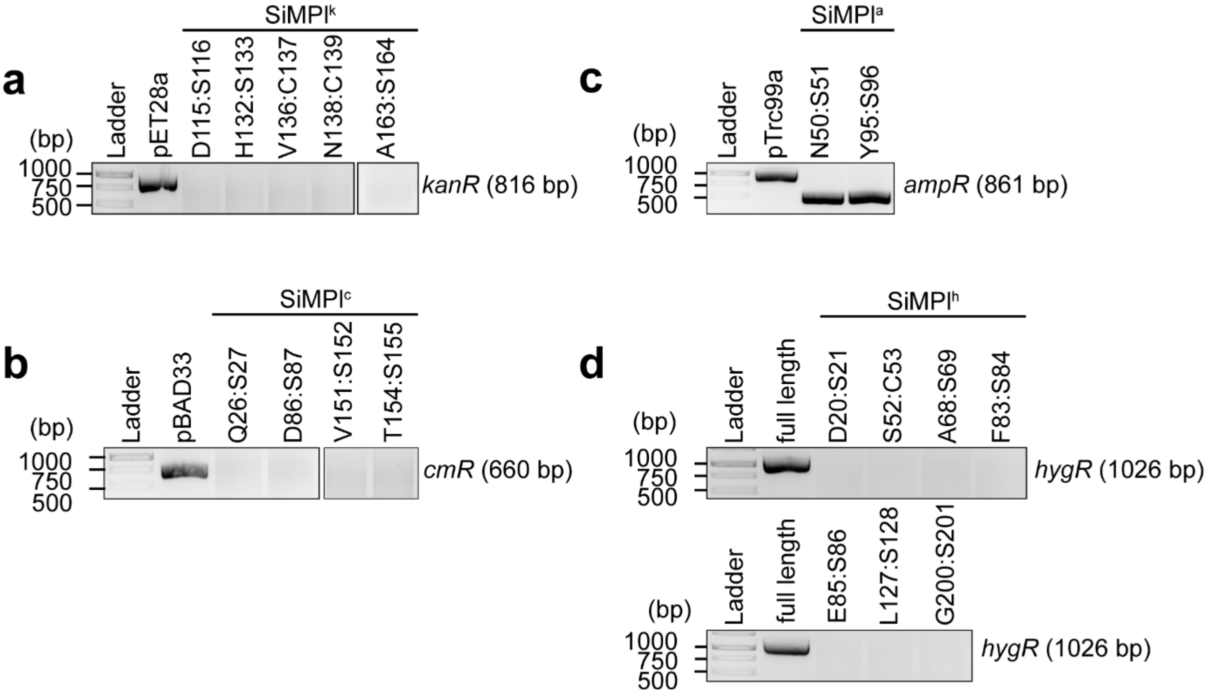


**Figure S7. A full-length resistance marker gene is not found in cells transformed with pSiMPl plasmid pairs.** (**a**-**d**) Representative ethidium bromide-stained 1% agarose gel showing the product of a PCR amplification with primers specific for the *kanR* (**a**), *cmR* (**b**), *ampR* (**c**) and *hygR* (**d**) genes, respectively. The PCR was performed on plasmid DNA extracted from cells transformed with the pSiMPl plasmid pairs or single plasmids encoding the full-length resistance genes, which were used as positive controls (pET28a, *kanR*; pBAD33, *cmR*; pTrc99a, *ampR*; pBAD33 into which the *cmR* gene was swapped with the *hygR* one, *hygR*). (**c**) pBAD33 carries a defunct *ampR* gene. So, you also see amplicons albeit smaller than the full-length gene.

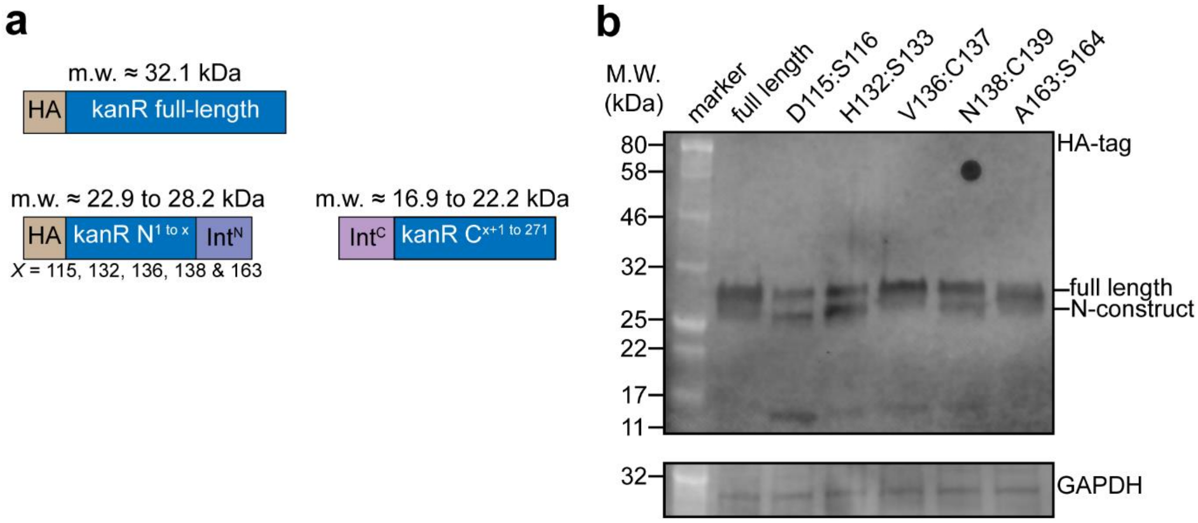


**Figure S8. Aminoglycoside 3′-phosphotransferase (APT) is reconstituted by gp41-1-mediated protein splicing.** (**a**) Schematics showing polypeptide fragments with predicted molecular weights. (**b**) Representative Western blot. The membrane was scanned with two different lasers to detect AlexaFluor 488-conjugated anti-rat secondary antibody directed against the rat anti-HA antibody (upper image) and AlexaFluor 790-conjugated anti-mouse secondary antibody directed against mouse anti-GAPDH (lower image).

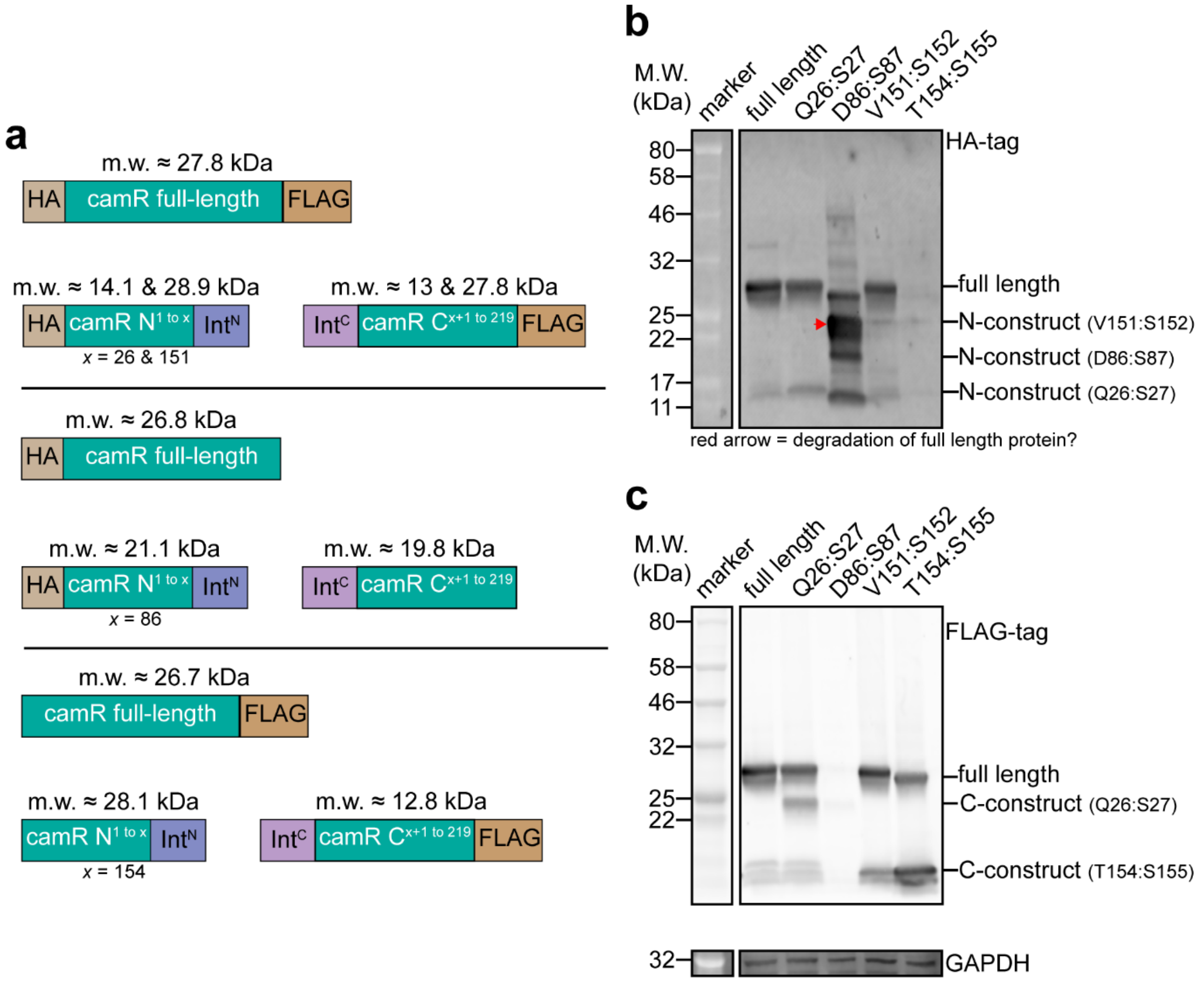


**Figure S9. Chloramphenicol acetyltransferase (CAT) is reconstituted by gp41-1-mediated protein splicing.** (**a**) Schematics showing polypeptide fragments with predicted molecular weights. (**b,c**) Representative Western blot. The membrane was scanned with three different lasers to detect AlexaFluor 488-conjugated anti-rat secondary antibody directed against the rat anti-HA antibody (upper image), Cyanine5-conjugated anti-rabbit secondary antibody directed against the rabbit anti-FLAG antibody (middle image), and AlexaFluor 790-conjugated anti-mouse secondary antibody directed against the mouse anti-GAPDH antibody (lower image).

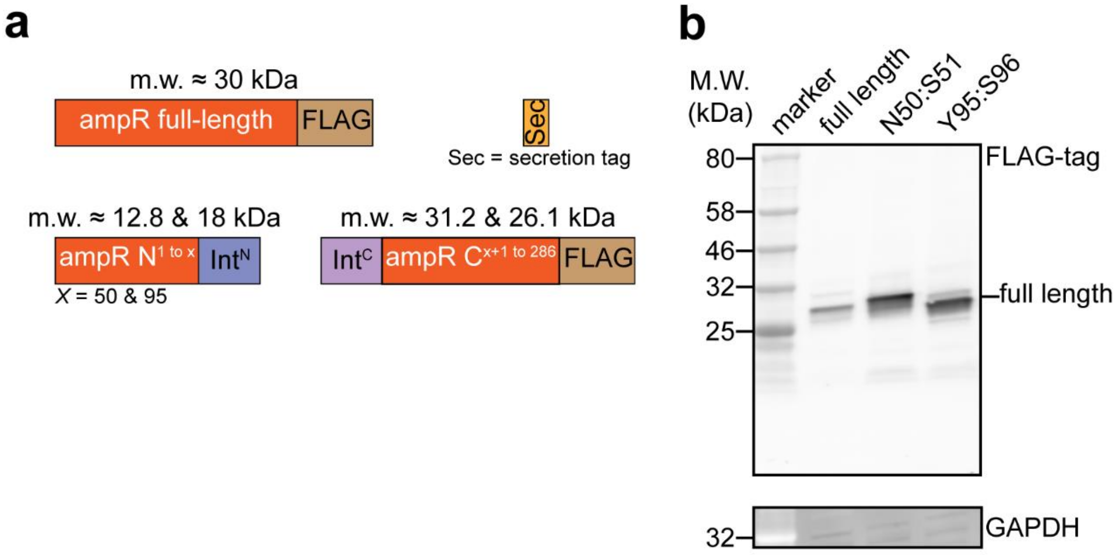


**Figure S10. TEM-1 β-lactamase (TEM-1 β-L) is reconstituted by gp41-1-mediated protein splicing.** (**a**) Schematics showing polypeptide fragments with predicted molecular weights. (**b**) Representative Western blot. The membrane was scanned with two different lasers to detect Cyanine5-conjugated anti-rabbit secondary antibody directed against the rabbit anti-FLAG antibody (upper image) and AlexaFluor 790-conjugated anti-mouse secondary antibody directed against mouse anti-GAPDH (lower image).

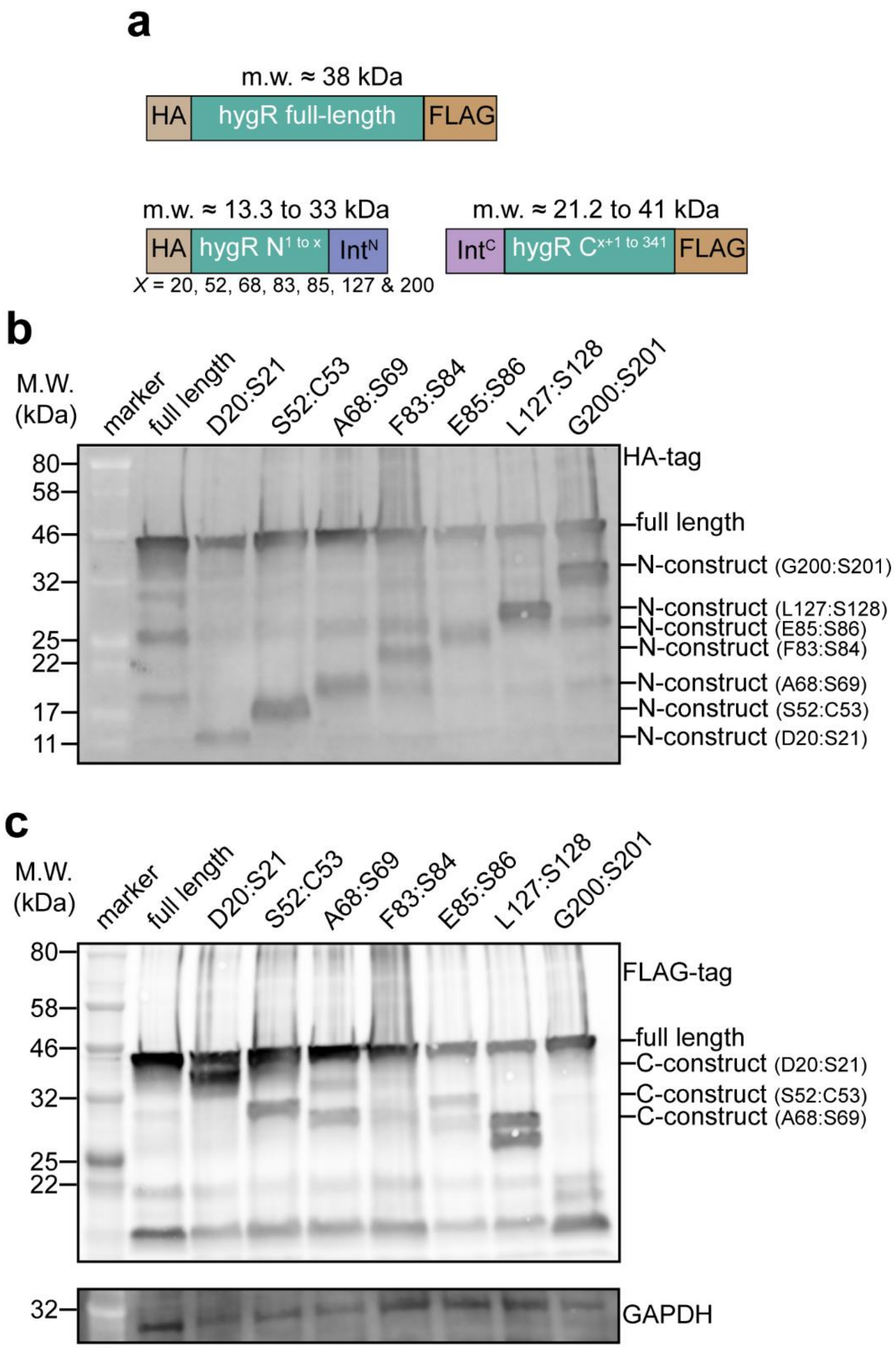


**Figure S11. Hygromycin B phosphotransferase (HPT) is reconstituted by gp41-1-mediated protein splicing.** (**a**) Schematics showing polypeptide fragments with predicted molecular weights. (**b,c**) Representative Western blot. The membrane was scanned with three different lasers to detect AlexaFluor 488-conjugated anti-rat secondary antibody directed against the rat anti-HA antibody (**b**), Cyanine5-conjugated anti-rabbit secondary antibody directed against the rabbit anti-FLAG antibody (**c**, upper image), and AlexaFluor 790-conjugated anti-mouse secondary antibody directed against the mouse anti-GAPDH antibody (**c**, lower image).

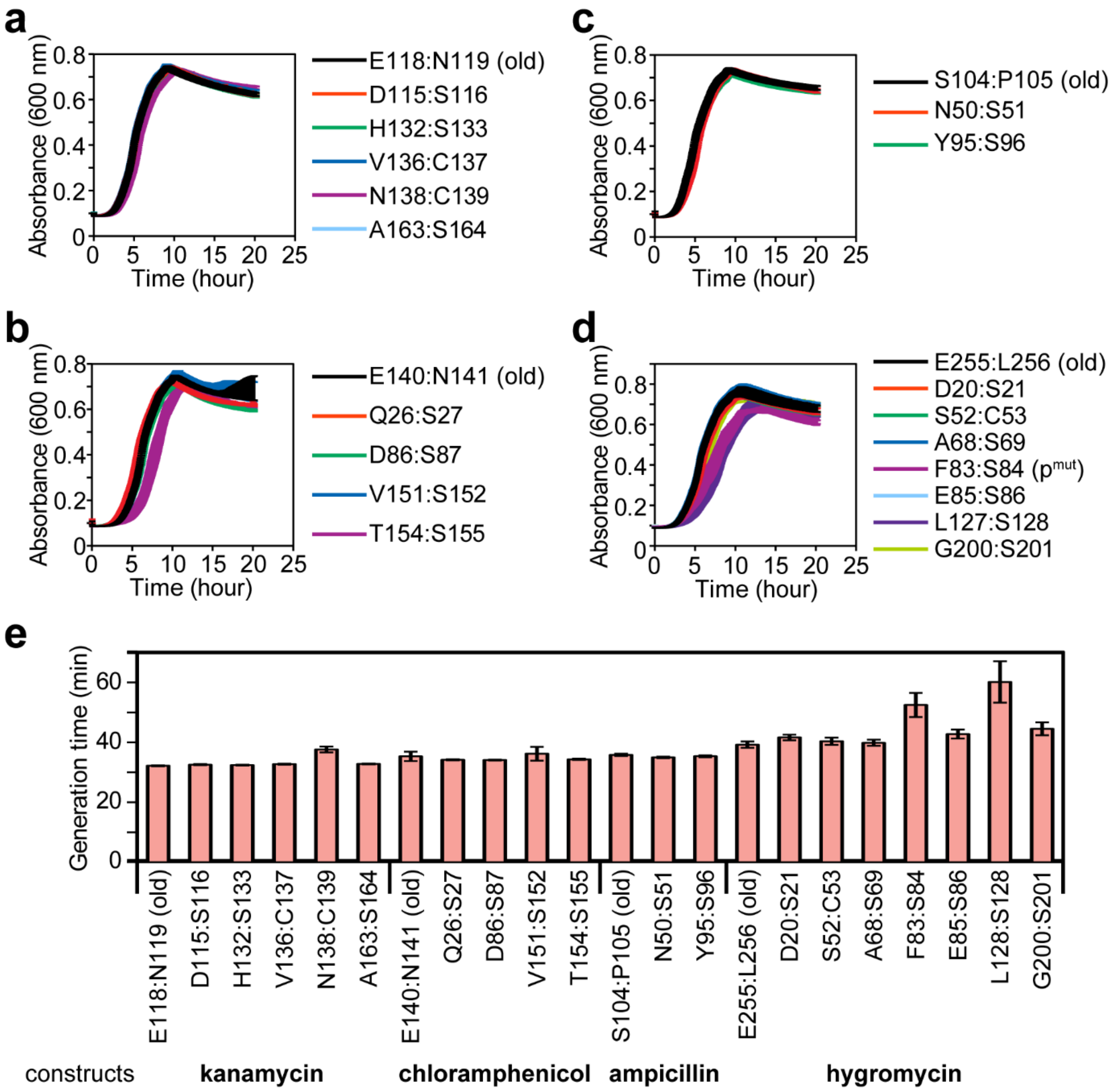


**Figure S12. Cells co-transformed with the pSiMPl plasmid pairs built splitting the enzymes at different sites grow similarly.** (**a-d**) Growth curves of *E. coli* TOP10 cells transformed with pSiMPl plasmid pairs for use with kanamycin (**a**), chloramphenicol (**b**), ampicillin (**c**), and hygromycin (**d**) built splitting the enzymes at the indicated sites. $p^{mut}$, mutated $P_{CAT}$ promoter. Old, previously identified splice site[2]. Experiments were performed twice (biological replicates) with technical triplicates each time. Values represent mean ± S.E.M. (**e**) Bar graph showing the generation time of bacteria grown in selective medium after transformation with the pSiMPl plasmid pair built splitting the enzymes at the indicated sites. The doubling time was calculated using data from (**a-d**) using the Growthcurver R package[3]. Values represent mean ± S.E.M..

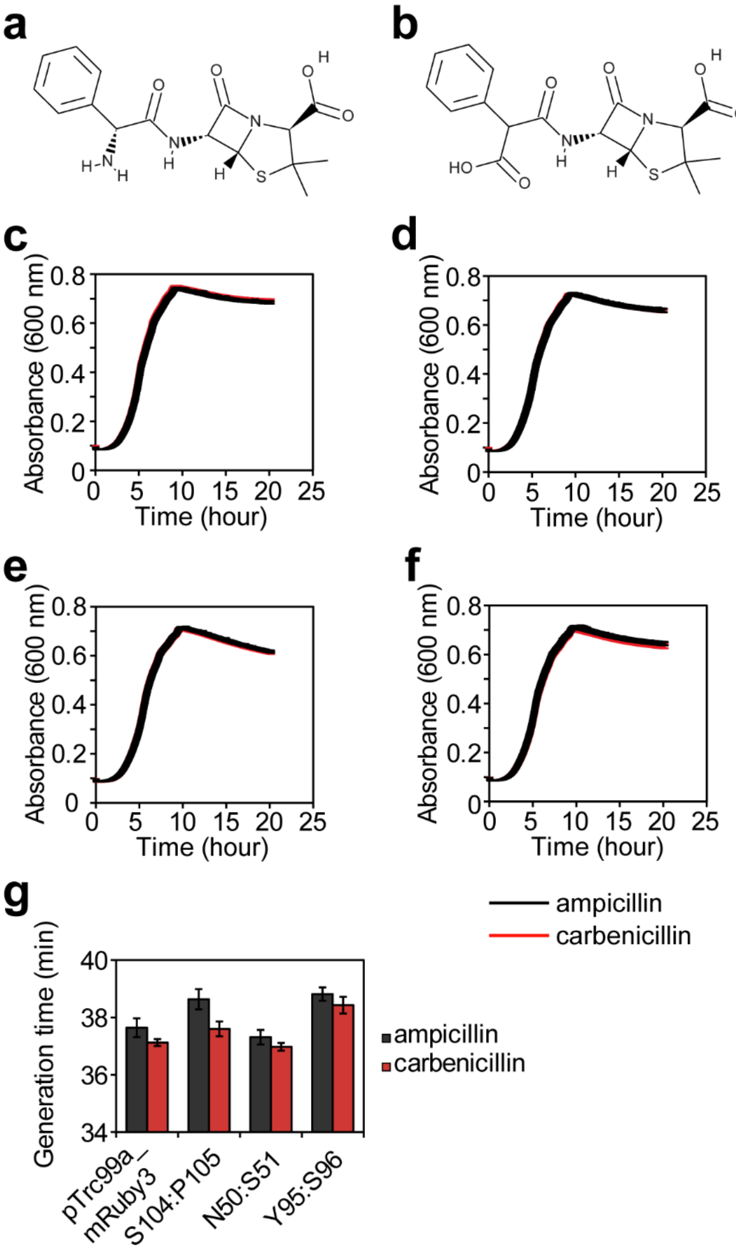


**Figure S13. Reconstituted TEM-1 β-lactamase expressed from pSiMPl[a]_N and pSiMPl[a]_C allows for growth on carbenicillin.** (**a**) Structure of ampicillin. (**b**) Structure of carbenicillin. (**c**-**f**) $OD_{600}$ measurements of *E. coli* TOP10 cells transformed with either pTrc99a-mRuby3 (**c**), pSiMPl[a] (splice site: S104:P105; old; **d**), pSiMPl[a] (splice site: N50:S51; **e**) and pSiMPl[a] (splice site: Y95:S96; **f**) grown in nutrient broth with the indicated antibiotic. Measurements were done in a plate reader for a total of 20 hours with each measurement taken every 2.4 min. Experiments were performed twice (biological replicates) with technical triplicates each time. Values represent mean ± S.E.M. (**g**) Generation time calculated using data from (**c**-**f**) with Growthcurver[3].

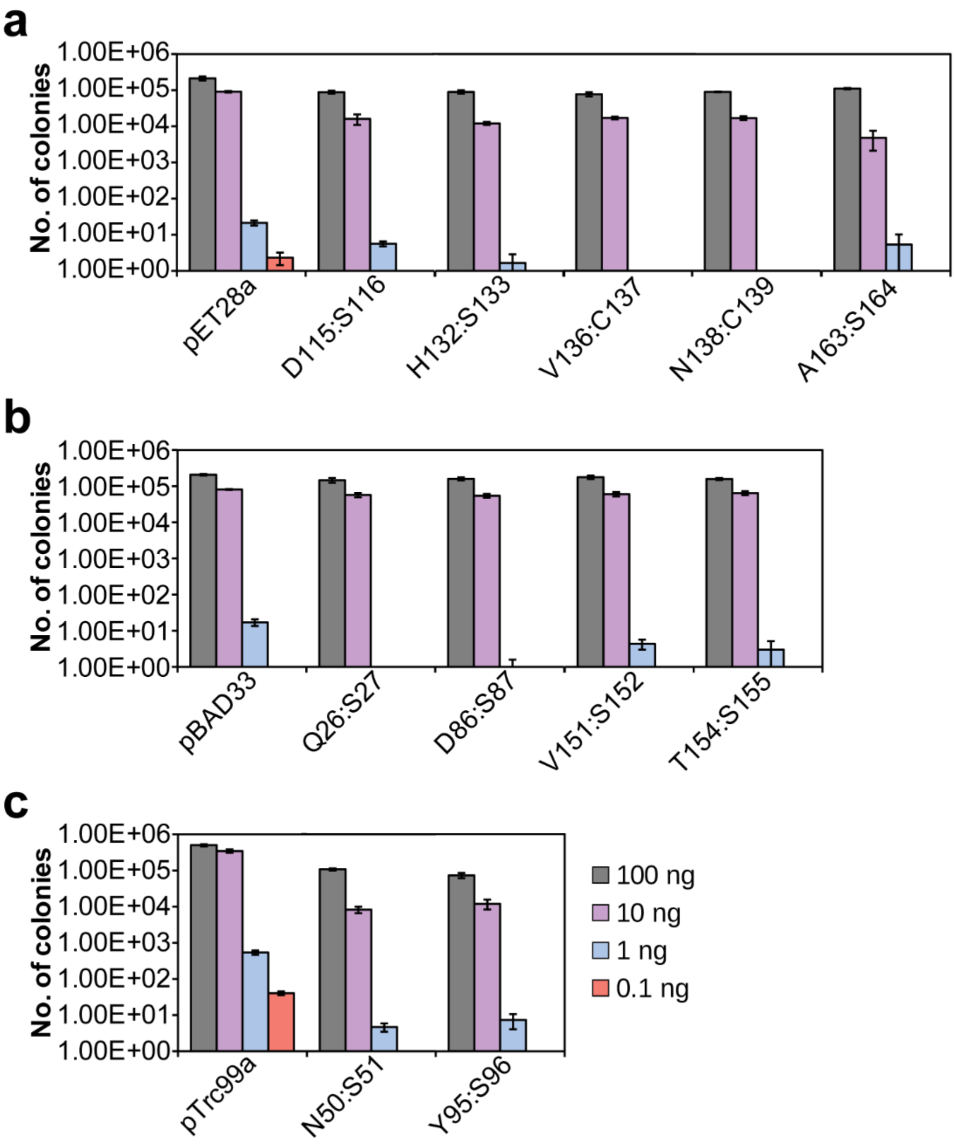


**Figure S14. Transformation efficiency of pSiMPl plasmids decreases with decreasing amounts of DNA.** (**a**-**c**) Bar graph showing the number of colonies obtained after transformation of chemically competent *E. coli* TOP10 cells with the indicated plasmids at the indicated amount of total plasmid concentration. Values represent mean ± S.E.M of three independent experiments. (**a**-**b**) color code as in (**c**). (**a**) kanamycin constructs, (**b**) chloramphenicol constructs and (**c**) ampicillin constructs. For the positive controls (pET28a, pBAD33 and pTrc99a) only one plasmid was transformed. In all other cases, the pSiMPl plasmid pair was co-transformed.

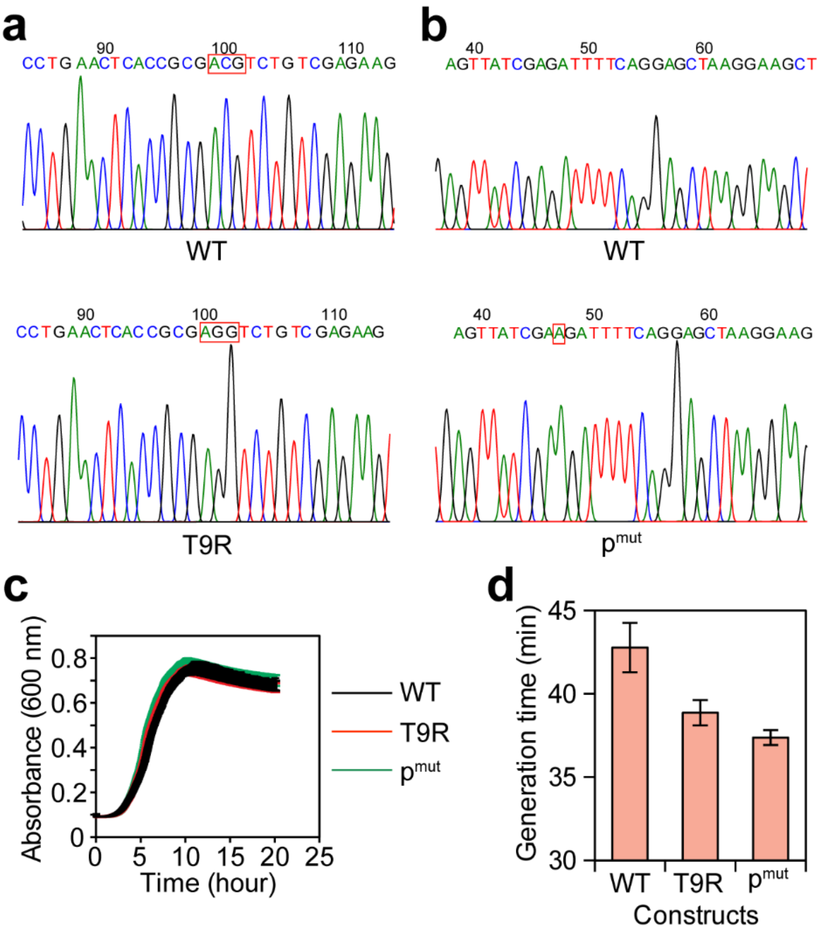


**Figure S15. Some pSiMPl$^h$_N constructs developed mutations.** (**a**-**b**) Representative chromatograms showing (**a**) the transversion mutation (C to G nucleotide change), which leads to the T9R mutation in the protein, and (**b**) the insertion mutation (insertion of nucleotide ‘A’ between potential transcription start site and ribosome binding site) identified in a few pSiMPl$^h$_N clones with the splice site: E85:S86. (**c**) Growth curve of *E. coli* TOP10 cells co-transformed with pSiMPl$^h$_C and either pSiMPl$^h$_N (splice site: E85:S86, with no mutations), pSiMPl$^h$_N with the transversion mutation (T9R) or pSiMPl$^h$_N with the insertion mutation ($p^{mut}$). $OD_{600}$ measurements were done in a plate reader for a total of 20 hours with each measurement taken every 2.4 min. Experiments were performed twice (biological replicates) with technical triplicates each time. Values represent mean ± S.E.M.. (**d**) Generation time calculated using data from (**c**) with Growthcurver[3].

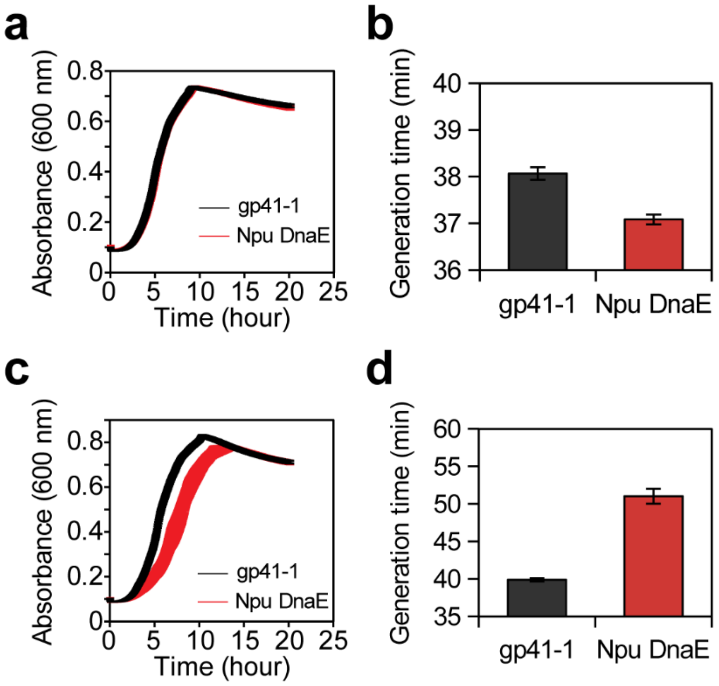


**Figure S16. pSiMPl plasmids can be constructed with different split inteins**. (**a**,**c**) $OD_{600}$ measurements of *E. coli* TOP10 cells transformed with $pSiMPl^k$ (splice site: V136:C137; **a**) and $pSiMPl^h$ (splice site: S52:C53; **c**) built using either the gp41-1 or *Npu* DnaE split intein. Measurements were done in a plate reader for a total of 20 hours with each measurement taken every 2.4 min. Experiments were performed twice (biological replicates) with technical triplicates each time. Values represent mean ± S.E.M.. (**b, d**) Generation time calculated using data from (**a**) and (**c**) with Growthcurver[3].

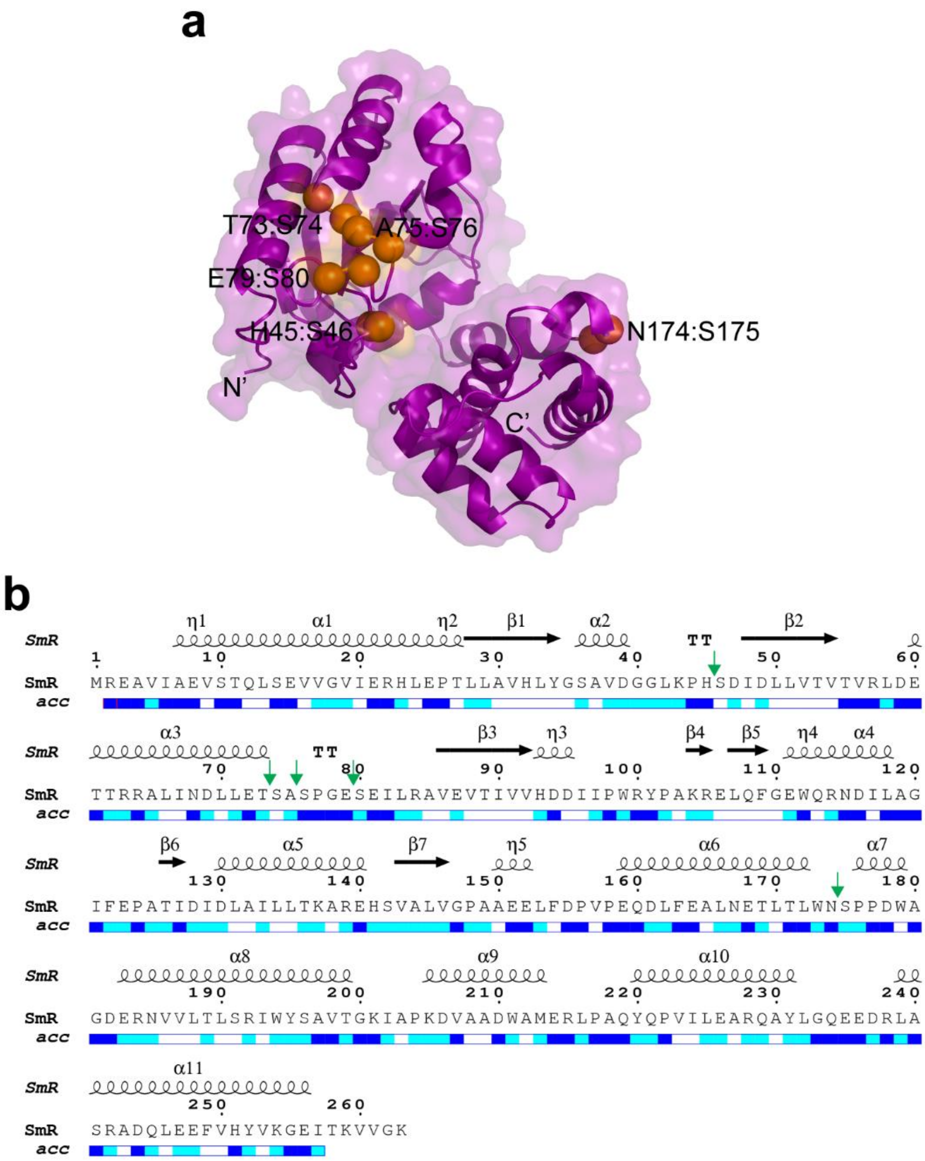


**Figure S17. The identified splice sites in aminoglycoside adenyltransferase (SmR) are surface accessible.** (**a**) Cartoon-surface representation of the structure of aminoglycoside adenyltransferase (SmR) modelled in SWISS-MODEL[4] using PDB ID: 5g4a as template. The residues at the splice site are indicated by orange spheres. Structures were depicted in PyMOL (PyMOL Molecular Graphics System, v. 1.8.x, Schrödinger, LLC). (**b**) Relative accessibility (acc) of each residue in aminoglycoside adenyltransferase. Secondary structural elements are represented by black arrows and coils. The image was created using the ESPript 3.0 server[1]. The color code is as follow: blue = accessible, cyan = partially accessible, white = buried, red = not predicted. Splice site are indicated by green arrows.

#### Materials and methods

##### Protein Cα fluctuation analysis

The X-ray structures of aminoglycoside 3′-phosphotransferase (PDB ID: 4ej7), chloramphenicol acetyltransferase (PDB ID: 1q23), TEM-1 β-lactamase (PDB ID: 1zg4) and hygromycin B phosphotransferase (PDB ID: 3w0s) were retrieved from the RCSB Protein Data Bank (https://www.rcsb.org/). Aminoglycoside adenylyltransferase structure was modelled on the SWISS-MODEL server[4] using the crystal structure of *Salmonella enterica* AadA as template (PDB ID: 5g4a) and SmR protein sequence from pCDF-1b as input sequence. The Cα fluctuation analysis was performed using the CABS-flex 2.0 webserver[5] with default parameters except for the seed value, which was changed in each independent coarse-grained simulation.

##### Plasmid construction

The $KanR^{1-118}$+SGY and SSS+$KanR^{119-271}$ regions from the previously constructed $pSiMPl^{k}$_N and $pSiMPl^{k}$_C plasmids[2] were removed by performing PCR separately using primer pairs Gib_SiMPl_BB1_FP and Gib_SiMPl_BB1_RP, and Gib_SiMPl_BB2_FP and Gib_SiMPl_BB2_RP, respectively. The complete list of primers used in the present study is given in **Supplementary File 1**. All PCR amplifications were performed with Phusion Flash High-Fidelity PCR Master Mix (2x) from ThermoScientific using Biometra-Thermocycler from analytik jena. The N- and C-terminal fragments of the resistance genes were amplified separately using the appropriate primer pairs (**Supplementary File 1**). These primer pairs were designed to have overlapping sequences at the 5' end with the respective plasmid backbones. Plasmids were finally assembled by Gibson Assembly®. Chemically competent *E. coli* TOP10 cells (Invitrogen) were used for all cloning purposes. For selection, kanamycin, chloramphenicol, ampicillin, hygromycin, spectinomycin and streptomycin were used at final concentrations 50 μg/mL, 35 μg/mL, 100 μg/mL, 100 μg/mL, 50 μg/mL and 50 μg/mL, respectively. Plasmid isolation was performed using the Miniprep kit from Qiagen. Sanger sequencing of the plasmids was done at GATC (Eurofins Genomics).

##### Bacterial growth analysis

Bacterial growth curves were acquired on the BioTek Synergy™ H4 plate reader as previously described[2]. Briefly, overnight *E. coli* TOP10 cultures carrying the plasmids were diluted to a starting $OD_{600}$ of 0.01 using fresh nutrient broth containing the necessary antibiotic. 120 μL of the freshly diluted cultures were transferred into a flat-bottom 96-well plate. The plate was sealed with the lid using parafilm tape. The parameters set were as follows: temperature = 37 $^{o}$C; run time = 20 h; read interval = 2 min and 43 s; wavelength = 600 nm; shake = slow; shake once every 130 s; read = absorbance end point; read speed = normal; delay = 100 msec. The plate reader was preheated to 37 $^{o}$C before the start of the experiment. Bacterial generation time (doubling time) was calculated using Growthcurver[3].

**Bacterial transformation efficiency analysis**

50 μL of chemically competent *E. coli* TOP10 cells (Invitrogen) were transformed with each SiMPl plasmid pair. Transformations were carried out by the standard heat shock method and either 100, 10, 1 or 0.1 ng of total plasmid DNA concentration were used for the transformation. As single plasmid controls, pET28a empty plasmid was used for kanamycin, pBAD33-EGFP was used for chloramphenicol and pTrc99a-mRuby3 was used for ampicillin. Colonies were counted manually.

**Western blot**

Western blots with bacterial samples were performed as previously described[2]. Bacterial cultures were grown from $OD_{600}$ = 0.1 until $OD_{600}$ ~1 at 37 °C 250 rpm, and 200 μL of each culture were centrifuged at 14800 r.p.m. for 1 min. Pellets were resuspended in 10 μL of 4x Laemmli sample buffer (Bio-Rad) and cells were lysed by incubating them at 95 $^{o}$C for 10 min. The samples were then loaded onto a 12% Mini-PROTEAN® TGX™ precast protein gel (Bio-Rad) and ran for ~1 hour at 120 V with 1x TGS running buffer. Proteins were transferred to PVDF membrane using Trans-Blot® Turbo™ Transfer Packs and a Trans-Blot® Turbo™ Transfer System (Bio-Rad) as per the manufacturer's protocol. After the transfer, membranes were first incubated with 4–5% BSA dissolved in TBST with shaking for 2 hours at room temperature, and rinsed with TBST. Then the membranes were incubated with either rat anti-HA (1:2000, Cat# 11867423001, Sigma), or rabbit anti-FLAG (1:2000, Cat# AHP1074, Bio-Rad), or mouse anti-GAPDH (1:1000, Cat# G13-61M, SignalChem) primary antibody at room temperature for 2 hours. Following 3 times wash (each time lasting for 5 min) with TBST, membranes were incubated with either AlexaFluor 488-conjugated anti-rat (1:2000, Cat# A-11006, Invitrogen), or Cyanine5-conjugated anti-rabbit (1:2000, Cat# A10523, Invitrogen), or AlexaFluor 790-conjugated anti-mouse (1:2000, Cat# A28182, Invitrogen) secondary antibody at room temperature for 1 hour. Unbound secondary antibodies were removed by washing the membrane twice with TBST (each time lasting for 5 min). Fluorescence signals were measured with an Amersham Typhoon imaging system (GE Healthcare) using 488, 635 and 785 nm wavelength lasers.
